# IGD: high-performance search for large-scale genomic interval datasets

**DOI:** 10.1101/2020.06.08.139758

**Authors:** Jianglin Feng, Nathan C. Sheffield

## Abstract

Databases of large-scale genome projects now contain thousands of genomic interval datasets. These data are a critical resource for understanding the function of DNA. However, our ability to examine and integrate interval data of this scale is limited. Here, we introduce the integrated genome database (IGD), a method and tool for searching genome interval datasets more than three orders of magnitude faster than existing approaches, while using only one hundredth of the memory. IGD uses a novel linear binning method that allows us to scale analysis to billions of genomic regions.

**Availability:** https://github.com/databio/IGD

## INTRODUCTION

Genome and epigenome data are frequently represented as *genomic intervals*, defined by a chromosome plus *start* and *end* coordinates. Because this data type is so common, interval-based comparisons are fundamental to genomic analysis^1–4^. The most basic operation on interval data is computing the intersections of two interval sets. This operation is common for a variety of use cases such as determining overlaps between sets of transcription factor binding sites, plotting the genome-wide distribution of a set of SNPs or regions, or assigning genomic annotations to query regulatory elements. Several approaches have been developed to do calculation this efficiently, such as the nested containment list (NCList)^5^, the interval tree^6^, the R-tree^7^, and the Augmented Interval List (AIList)^8^, and many tools exploiting these data structures exist^1,9–12^.

A more advanced task with intervals is *interval database search*, which requires comparing a query interval set against a database of multiple interval sets. This task extends the file-to-file comparison to a file-to-database comparison. To do this, the file-to-file approaches can be applied in a loop, but this does not take advantage of integrating across interval sets in the database, which can be done by creating a combined index that spans individual interval sets. This is the approach taken by GIGGLE, which is much faster for searching a large database than a naive file-to-file loop^3^. Here, we present a new approach called the *integrated genome database (IGD)*, which uses a novel indexing method called *linear binning*. We show that the IGD approach achieves substantially better performance than all existing approaches.

## METHODS

### Linear binning

An interval set has two different orders: the order of start values and the order of end values. To effectively process interval data, multi-level binning methods are usually used^2,7,10^, where intervals are stored in bins of different sizes that are large enough to contain the intervals. In contrast, the linear binning approach used by IGD is a single-level binning (Fig. 1a). The whole genome is divided into equal-sized bins and intervals are put into bins they intersect. The intervals are not required to fit into the bins; instead, intervals that cross bin boundaries are stored in multiple bins. This duplication introduces two challenges: increase in size of the database, and a possibility of multiple counting. The number of duplicates can be adjusted by the selecting bin size, which therefore adjusts the amount of extra data required for linear binning. In our tests, the database size increases by only 5% to 20% for a bin size of 16,384 (2^14^), which is much more efficient than multi-level binning or tree-based methods (see results in the next section). The multiple counting challenge, which occurs only when both the query and the data interval cross the same bin boundary, can be neatly solved by setting a rule: if the query crosses the left boundary of the bin, then any region in the bin that also crosses the left boundary will be skipped. In figure 1a, query *q*2 intersects both interval 2*a* (which belongs to bin *i*) and interval 2*b* (bin *i* + 1); since *q*2 does not cross the left boundary of bin *i*, region 2*a* will count; but *q*2 crosses the left boundary of bin *i* + 1, so the intersection with region 2*b* will not count.

**Fig. 1:**
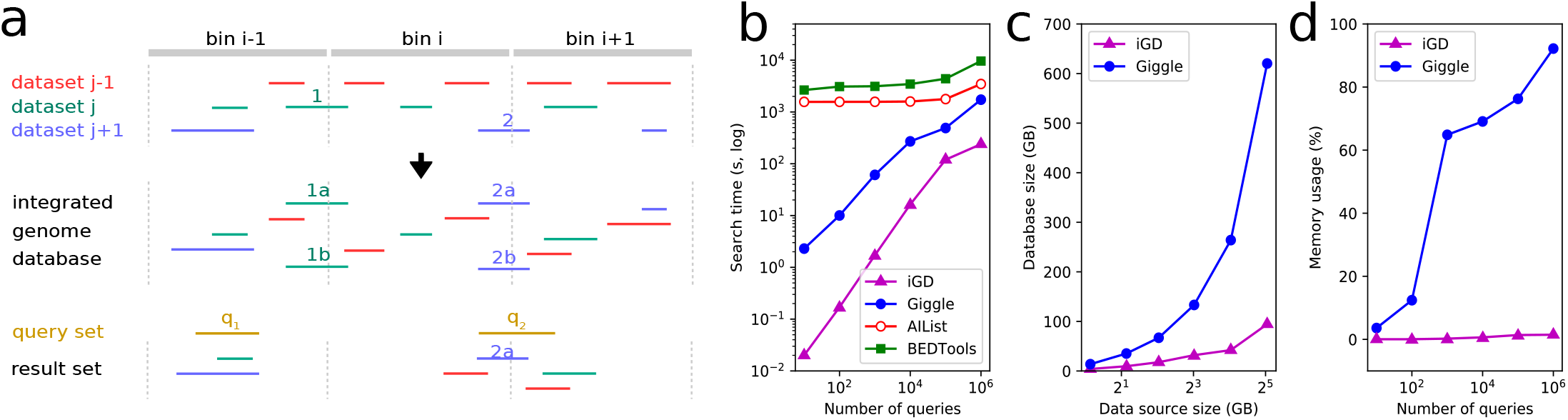
a) Linear-binning indexing. The genome is divided into equal-sized bins and regions are put into any bins they intersect. Regions that cross boundaries are stored in multiple bins. For example, region 1 is placed in both bin i − 1 and bin i. Regions in each bin are then sorted by their starts. Search queries may be unsorted. Colors indicate that the file each region belongs to. A search query interval q2 crosses the boundary of bin i and bin i + 1, so region 2 is double counted. IGD solves this by counting only 2a (see text); b) Performance comparisons with existing methods on 5660 interval datasets. Search times are shown for IGD, GIGGLE, AIList and BEDTools on 6 query datasets of 10^1^ to 10^6^ regions; c) Sizes of IGD and GIGGLE databases created from 6 different subsets of a UCSC interval set collection with input size ranging from 2^0^ to 2^5^ GB of compressed data; d) Peak memory usage for IGD and GIGGLE for the 6 queries. GIGGLE did not complete the query of 10^6^ regions.

With the challenges of multiple counting and database size neatly solved, linear binning makes interval data behave similar to a dataset with a single order. More importantly, since a bin contains all regions that intersect the bin, the processing of intervals in each bin is independent. In contrast, the nodes of the tree-based or multi-level approaches are not independent and queries typically require traversing nodes or bins of different levels.

### IGD data structure and query

IGD stores the bin data linearly and maintains a header that contains the location of the bins and the number of IGD data elements in each bin. Each IGD data element contains the index of the original dataset, the original genomic region (*start* and *end* coordinates), and a value (signal level). For compatibility to other existing search tools, IGD also supports a data element type without the signal value. Since the region data is indexed linearly, a query to the database simply determines the index of the bin that the query region intersects and then loads only that bin into memory and searches within the bin. If a query crosses any bin boundaries, IGD loads and searches those bins one by one. Compared to file-to-file approaches, IGD does not need to load all the files and does not need to search all regions in every file, making IGD much more efficient (Fig. 1b).

Regions in a bin are presorted by the region *start*, so the search inside a bin can be done by first using a binary search to find the right-most interval (intervals with *start* larger than or equal to the query *end* will not intersect the query) then scanning to the left (smaller *start*). This is very efficient because the bin size, *e.g.*, 16,384 bp, is small. Alternatively, the general search algorithm of AIList^8^ may be used, but it requires preprocessing to add an extra element (the augmenting value) for each region, which increases the storage by one third and also affects the data loading speed.

## RESULTS

To benchmark the speed and memory use of the IGD approach, we assembled a test set of human query and database interval sets. All tests in this work were performed using a single core on a 2.8 GHz personal computer with 16 GB memory and an external 1TB SSD disc. We created the IGD databases with a bin size of 8,192 (2^13^), corresponding to about 376,000 bins for human genome.

We first compared IGD to existing approaches for the database search task. For query sets with 100 or fewer intervals, IGD is four to five orders of magnitudes faster than file-to-file approaches AIList and BEDTools (Fig 1b). This is expected, as these approaches load all data from 5660 files while IGD only loads data from very few bins. GIGGLE uses a database approach based on B+ tree indexing scheme, which is faster than the file-by-file methods, but IGD is two orders of magnitude faster still. For larger query intervals, IGD is also several orders of magnitude faster than any existing approach.

In terms of disk use, the IGD index increases linearly with the size of input data source, while GIGGLE increases exponentially (Fig 1c). For the full set UCSC data source, GIGGLE uses 640GB disc space while IGD only needs 98GB. IGD is also faster in database creation: the complete database takes IGD about 2 hours to create, while GIGGLE takes more than 4 hours.

Furthermore, IGD uses nearly negligible computer memory while GIGGLE needs substantial memory even for small query sets (Fig. 1d). For example, with 1,000 query regions, IGD requires only 38 MB of memory, while GIGGLE uses 10,570 MB memory, more than 270 times as much. These results suggest that linear-binning indexing is superior to B+tree indexing in terms of database creation time, search time, and search memory use. Bin size is an option set by the user when a database is created. To explore how bin size affects search speed and duplication rate, we compared various datasets with bin sizes ranging from 2^12^ to 2^18^ and found that the default bin size (2^14^) is near optimal for all scenarios we tested (Supplementary Information). We conclude that for most use cases, the user will not need to worry about bin size.

We have shown how IGD presents a fast alternative for interval search operations. Furthermore, the independence of IGD bins and the small memory use make IGD amenable to parallel processing, which can further increase the searching speed. In additional to calculating number of overlaps, IGD can also calculate the seqpare metric, which provides a novel way to assess similarity between region sets^13^. IGD is well suited for tasks that require searching a database of intervals with a query, such as in region set enrichment overlap calculations^12,14,15^, or methods that use collections of region sets to aggregate data^16–19^. As the size of genomic interval data increases and more databases arise to serve and analyze this data type^14,20–22^,we believe the speed and memory advances achieved by IGD will make it a valuable tool for fast analysis in the future.

## Supplementary Information

### 1. The impact of bin size on database size and search speed

The choice of the bin-size (*l*_*bin*_) for IGD has greater impact on the size of the created database than on the search speed. Theoretically, this impact depends on the average interval length (*l*_*avg*_) of the interval sets involved. The rate of bins that split the intervals (the duplication rate) is *l*_*avg*_/*l*_*bin*_. Therefore, if *l*_*bin*_ is close to *l*_*avg*_, the size of the created IGD database will be twice of the size of the original datasets; to make IGD database size 1/4 larger than the original size, *l*_*bin*_ should be 4 × *l*_*avg*_. Since IGD uses binary search to determine the overlaps in a bin, 4 × changes of *l*_*bin*_ only add 2 extra search operations, which has only a minor effect on overall speed.

The following two tables list test results on the database size and search speed for 3 representative collections of interval sets with a wide range of average lengths: roadmap (*l*_*avg*_=7064), a subset of UCSC (*l*_*avg*_=1290), and a subset of JASPAR (*l*_*avg*_=9).

**Table S1.**
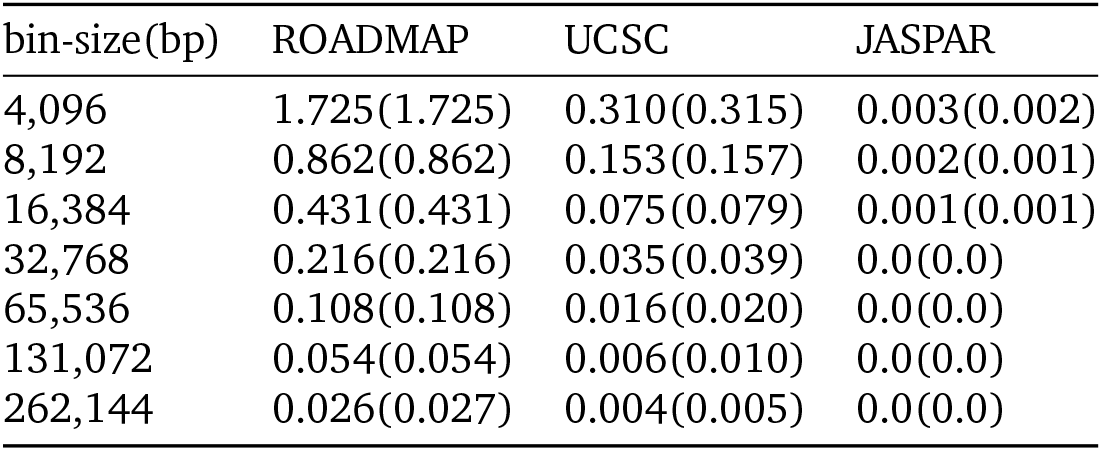
Measured duplication ratio ((IGD database size-original size)/original size) of IGD database at different bin-sizes for three different data sources. The calculated duplication ratios are listed in parentheses. For ROADMAP data, the measured values are nearly exact as calculated (*l*_*avg*_/*l*_*bin*_).

**Table S2.** Runtime(s) of IGD search for databases created using different bin-sizes (Table S1). The query interval set contains 1 million intervals.

| bin-size(bp) | ROADMAP | UCSC | JASPAR |
| --- | --- | --- | --- |
| 4,096 | 8.372 | 20.126 | 8.668 |
| 8,192 | 5.123 | 15.556 | 7.884 |
| 16,384 | 3.806 | 12.975 | 7.522 |
| 32,768 | 3.239 | 12.906 | 7.290 |
| 65,536 | 3.055 | 13.664 | 7.478 |
| 131,072 | 2.890 | 15.251 | 7.439 |
| 262,144 | 3.125 | 18.989 | 10.34 |

It can be seen that bin-size of 16384, 32768 and 65536 are generally good for search for databases with regions that range in average size from 9 to 7000+. The default bin-size is set to 16384 (2^14).

### 2. Performance of IGD on different computing enviroments

The current implementation of IGD is serial, although it has great potential for parallel processing. We tested IGD on both a personal computer and on a high performance cluster (HPC) compute node. IGD runs nearly identically in either case, assuming the hard disk type is the same. With a conventional hard-disk-drive (HDD), results will be similar whether on and on a node of a local cluster when the cluster node has the same processing power as the personal computer. However, we found that IGD runs substantially faster when the database is stored on a solid-state-drive (SSD). With an SSD, the IGD database creation is 3 to 6 times faster and IGD database search is roughly 2 times faster. Therefore, to create an IGD database from large-scale datasets like UCSC (^~^100GB), a large volume SSD (1TB or larger) is recommended.

